# uniPort: a unified computational framework for single-cell data integration with optimal transport

**DOI:** 10.1101/2022.02.14.480323

**Authors:** Kai Cao, Qiyu Gong, Yiguang Hong, Lin Wan

## Abstract

Single-cell data integration can provide a comprehensive molecular view of cells. Here we introduce uniPort, a unified single-cell data integration framework which combines a coupled Variational Autoencoder (coupled-VAE) and Minibatch Unbalanced Optimal Transport (Minibatch-UOT). It leverages both highly variable common and dataset-specific genes for integration and is scalable to large-scale and partially overlapping datasets. uniPort jointly embeds heterogeneous single-cell multi-omics datasets into a shared latent space. It can further construct a reference atlas for online prediction across datasets. Meanwhile, uniPort provides a flexible label transfer framework to deconvolute spatial heterogeneous data using optimal transport space, instead of embedding latent space. We demonstrate the capability of uniPort by integrating a variety of datasets, including single-cell transcriptomics, chromatin accessibility and spatially resolved transcriptome data. uniPort software is available at https://github.com/caokai1073/uniPort.

## Introduction

The latest developments in high-throughput single-cell multi-omics sequencing technologies, e.g., single-cell RNA-sequencing (scRNA-seq) and ATAC-sequencing (scATAC-seq), enable comprehensive studies of heterogeneous cell populations that make up tissues, the dynamics of developmental processes, and the underlying regulatory mechanisms that control cellular functions. The computational integration of single-cell datasets is drawing heavy attention toward making advancements in machine learning and data science [1,2]. Among existing single-cell integration methods, tremendous efforts have been devoted to integrating multiple datasets simultaneously profiled from the same cells [3–5] (e.g., paired-cell datasets generated by the cellular indexing of transcriptomes and epitopes by sequencing (CITE-seq) [6]). However, these paired datasets are technically challenging and costly to obtain. Therefore, a vast number of integrative methods have been developed for data profiled from different cells taken from the same, or similar, populations. For example, the celebrated platform Seurat [7] projected feature space into a common subspace using canonical correlation analysis (CCA), which maximizes inter-dataset correlation. LIGER [8] and DC3 [9] employed non-negative matrix factorization to find the shared low-dimension factors of the common features to match single-cell omics datasets. Harmony [10] iterated between maximum diversity clustering and a mixture model-based linear batch correction to simultaneously account for multiple experimental and biological factors. However, these methods rely on linear mapping, thus lacking the ability to handle nonlinear deformations across cellular modalities. In addition, they only leverage filtered common genes, while ignoring the importance of dataset-specific genes for identification of cell populations, which usually capture cell-type heterogeneity not present in the common genes. To address these shortcomings, manifold alignment methods are emerging and have achieved promising results in integrating single-cell multi-omics datasets [11–14]. However, understanding of the properties of latent biological manifolds is incomplete, and alignment of latent manifolds cannot be guaranteed [2]. Furthermore, manifold alignment methods are limited by relatively high computational complexity, and they are not scalable to large-scale datasets.

With the development of deep learning, many neural-network approaches have been proposed and demonstrated powerful in data integration across modalities. Most current neural-network methods are based on autoencoder and require paired-cell datasets, such as DCCA [15] and Cobolt [16], to utilize cell-paring information. However, when cell-paring information is unavailable, simultaneously training different autoencoders and aligning cells across different modalities in latent space still make computation challenging exercise. Recently, an emerging number of neural-network methods have been developed account for unpaired data. For example, methods like scDART [17] and cross-modal autoencoders [18] attempt to combine distribution alignment methods with autoencoder, e.g., Maximum Mean Discrepancy (MMD) or a discriminator network. However, these distribution alignment terms require distributions across modalities to be matched globally, which is often too restrictive when datasets partially overlap. SCALEX [19] resolves the limitation raised by global matching by simply excluding an explicit alignment constraint, but it also restricts its ability to integrate heterogeneous datasets. SCALEX only utilizes common genes, such as those used by linear mapping methods. In addition, the transfer learning-based methods were also developed to establish a source atlas via one modality for knowledge (e.g., cell labels) transfer to another modality by learning a modality-invariant latent space [20,21]. Although having achieved encouraging results, these methods are restricted to using source modality with cell label annotated.

Here we introduce uniPort, a unified deep learning method that integrates heterogeneous singlecell datasets. It leverages cellular low-dimensional common space and cell heterogeneity by combining a coupled Variational Autoencoder (coupled-VAE) and Minibatch Unbalanced Optimal Transport (Minibatch-UOT) [22]. Our uniPort efficiently integrates diverse and heterogeneous single-cell datasets, e.g., single-cell transcriptomics, chromatin accessibility and spatially resolved transcriptome data. uniPort can not only jointly embed heterogeneous datasets into a latent space, but also construct a reference atlas for online prediction across modalities when only one modality is present. Meanwhile, uniPort can deconvolute compounded data in a broader application spectrum of resolution maps, such as spatially resolved high-plex RNA imaging-based data and barcoding-based spatial transcriptome (ST) data. Concretely, uniPort (1) leverages both highly variable common and dataset-specific genes for integration in a manner that allows the integrated latent space to be further revised throughout training; (2) employs a novel coupled-VAE neural network modal that strengthens the power of nonlinear correction and leverages the generalization ability of coupled-VAE to construct a reference atlas for online prediction across modalities; (3) minimizes a Minibatch-UOT loss, which is scalable to large datasets and suitable for partially overlapping alignment, providing a strong guarantee for heterogeneous data integration and removing the constraint of paired cells such as that found in other autoencoder-based modals; and, finally, (4) outputs a global optimal transport (OT) plan, i.e., a cell-cell correspondence matrix, that provides flexible transfer learning for deconvolution of spatial heterogeneous data using OT space, instead of embedding latent space. Experimental results show that uniPort outperforms other methods in integrating datasets profiled from peripheral blood mononuclear cells (PBMC). It is also a powerful online predictor of spatially resolved data through scRNA-seq data by a well-trained atlas. Moreover, with a global OT plan, we demonstrate that uniPort can successfully decipher canonical structures of mouse brain and assist in locating Tertiary lymphoid structures (TLS) in the breast cancer region, as well as reveal cancer heterogeneity in microarray-based spatial data.

## Results

### uniPort embeds and integrates datasets by coupled-VAE and Minibatch-UOT

As input, uniPort takes diverse and heterogeneous single-cell datasets across different modalities or technologies. uniPort is based on a coupled Variational Auto-Encoder (VAE) and first employs a dataset-free encoder to project highly variable common gene sets of different datasets into a generalized cell-embedding latent space. Then uniPort reconstructs two terms. One is input by a dataset-free decoder with Dataset-Specific Batch Normalization (DSBN) [19,23] layers. The other is a highly variable gene set through a dataset-specific decoder corresponding to each dataset (Fig. 1 and Supplementary Fig. 1). Some overlapping genes are often found between the two terms as some common genes are also highly variable in each dataset. However, with slight abuse of ‘specific’, we still name the second term a ‘dataset-specific’ gene set in the following context. During integration, uniPort minimizes a Minibatch-UOT loss between cell embeddings in the latent space from different datasets. It is necessary to introduce the loss as it feeds back a gradient to the encoder to achieve a better alignment result, especially when dataset-specific decoders are considered that increase the heterogeneity across datasets in the latent space. Meanwhile, the mini-batch strategy substantially improves the computational efficiency of optimal transport, making it scalable to large datasets, and the unbalanced mode makes it more suitable for partially overlapping integration. We adopt an efficient alternate optimization method to solve the problem (Methods).

**Figure 1:**
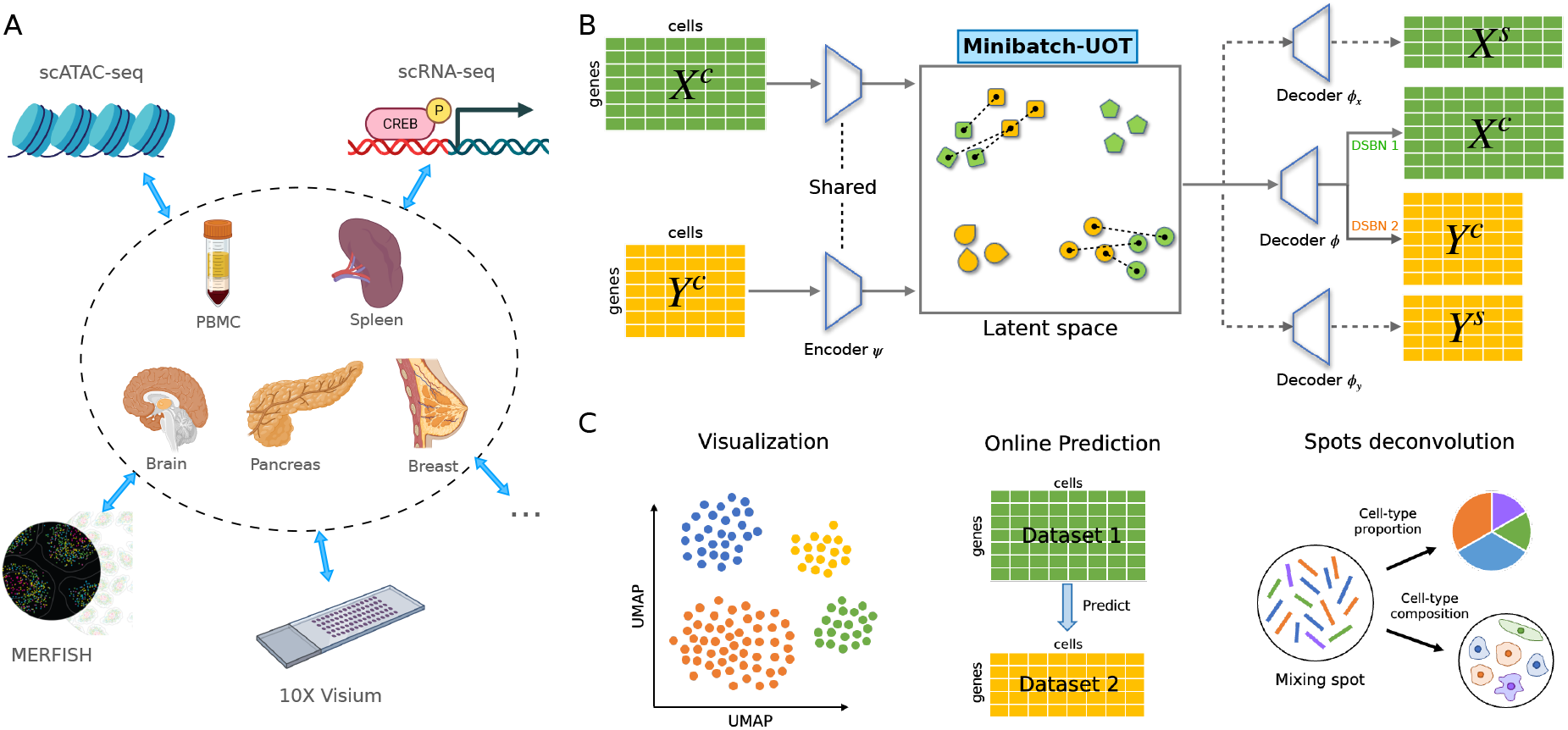
Overview of uniPort algorithm. uniPort integrates single-cell data by combining a coupled-VAE and Minibatch-UOT. **(A)** uniPort takes as input a highly variable common gene set of single-cell datasets across different modalities or technologies. **(B)** uniPort projects input datasets into a cell-embedding latent space through a shared probabilistic encoder. Then uniPort minimizes a Minibatch-UOT loss between cell embeddings across different datasets. Finally, uniPort reconstructs two terms. The first consists of input datasets by a decoder with different DSBN layers. The second consists of highly variable gene sets corresponding to each dataset by dataset-specific decoders. **(C)** uniPort outputs a shared latent space and a global optimal transport plan that can be used for downstream analysis, such as visualization, online prediction and spots deconvolution.

We compared uniPort to current state-of-the-art single-cell data integration methods, including Seurat v3, LIGER, Harmony, MultiMAP [24] and SCALEX. For this purpose, we examined two scATAC and scRNA datasets, including peripheral blood mononuclear cells (PBMC) from 10X Genomics and mouse spleen [24, 25] from MultiMAP (Supplementary Fig. 2). We also considered the integration of heterogeneous spatial transcriptome (ST) with scRNA data. Two main types of ST sequencing technologies are high-plex RNA imaging-based and barcoding-based. High-plex RNA imaging-based spatial sequencing has the advantage of single-cell precision with greater depth, but it is restricted to partial measurement with lower coverage. To test the performance of uniPort over the high-plex RNA imaging-based data, we applied uniPort to integrate the scRNA data and the spatially resolved multiplexed error robust fluorescence in situ hybridization (MERFISH) data [26]. Barcoding-based ST, in contrast, is more accessible to transcripts and achieves higher coverage, but it is limited to the mixing spots with lower resolution [27]. On the other hand, uniPort is able to transfer labels by a global OT plan, instead of training a k-nearest-neighbor (kNN) classifier in the latent space, which is more suitable for deconvolution of compounded barcoding-based ST data in a broader application spectrum of resolution maps. Therefore, we integrated (1) ST data from 10X Visium, including mouse brain [28] and HER2-positive breast cancer tissues [29], and (2) low-resolution microarray-based ST data from pancreatic ductal adenocarcinoma tissues [30], with their matched scRNA data, separately.

We employed a series of scores to assess the performance of single-cell data integration. To quantify dataset mixing and cell-type separation, we computed two scores used by SCALEX: the Batch Entropy score [31] to evaluate the extent of mixing cells across datasets, and the Silhouette score [32] to evaluate the separation of biological distinctions. To benchmark annotation clustering accuracy, we adopted the adjusted Rand Index (ARI) [33], the Normalized Mutual Information (NMI) and the F1 scores using cell-type annotations.

### uniPort integrates single-cell transcriptomics and chromatin accessibility

We first applied uniPort to integrate transcriptomic and epigenomic data using scATAC and scRNA datasets profiled from peripheral blood mononuclear cells (PBMC) (Fig. 2), including 11259 paired cells with 19434 features in scATAC and 26187 genes in scRNA. We employed Uniform Manifold Approximation and Projection (UMAP) [34, 35] to visualize the integration performance of all methods separately (Fig. 2A, Supplementary Fig. 3). We also involved a uniPort version that only used common genes for integration, named uniPort_cm for convenience.

**Figure 2:**
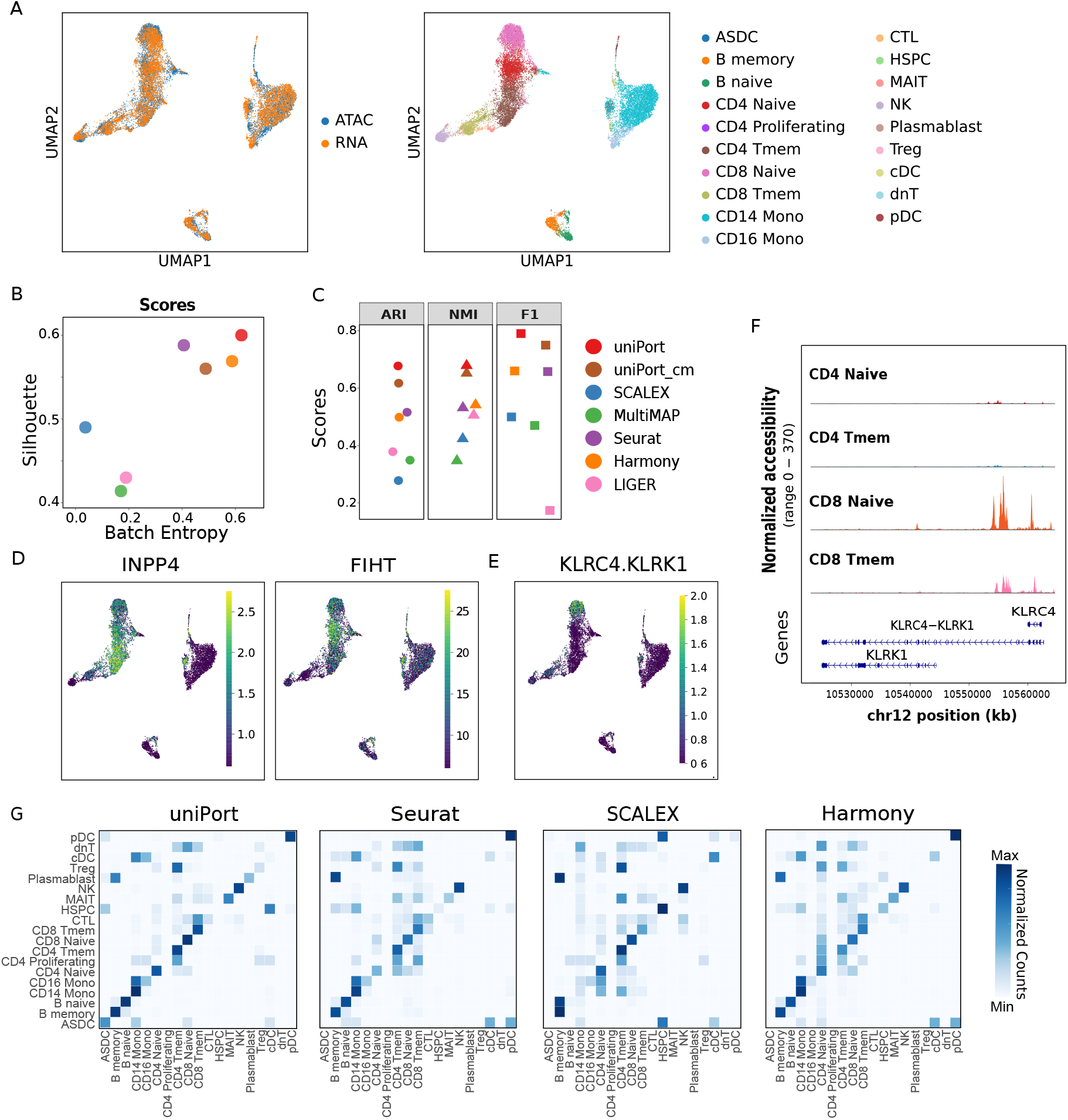
uniPort integrates single-cell transcriptomics and chromatin accessibility. **(A)** UMAP visualization of PBMC data after integration colored by omics and cell annotations. **(B)** Comparison of Batch Entropy and Silhouette scores for evaluating integration performance. **(C)** Comparison of ARI, NMI and F1 scores for evaluating clustering performance. **(D)** UMAP visualization of expression levels of RNA-specific marker genes INPP4 and FIHT. **(E)** UMAP visualization of expression levels of ATAC-specific marker gene KLRC4.KLRK1. **(F)** DNA accessibility of KLRC4.KLRK1 loci. **(G)** Comparison of confusion matrices of label transfer from scRNA-seq data to scATAC-seq data.

uniPort effectively integrated the scATAC and scRNA data by common, RNA-specific and ATAC-specific (in the form of gene activity matrix) highly variable genes without using pairing information. We evaluated the performance of data integration via the Silhouette and the Batch Entropy scores, as well as the widely used ARI, NMI and F1 scores using cell-type annotations, as noted above (Fig. 2B,C). Overall, uniPort outperformed all other methods and achieved the highest scores, including Silhouette score of 0.6, Batch Entropy score of 0.622, ARI of 0.677, NMI of 0.679 and F1 of 0.79. In comparison, uniPort_cm, with only common genes, achieved the second highest ARI of 0.617, NMI of 0.651 and F1 of 0.749, and Harmony ranked second in Silhouette score (0.56) and Batch Entropy score (0.488). In contrast, SCALEX, MultiMAP and LIGER did not perform well in this task since all evaluated scores were below 0.5.

As visualized by UMAP, uniPort, uniPort_cm, Seurat and Harmony accurately integrated most cell types in two modalities, including CD8 Naive, CD4 Naive, CD14 Mono and B cells (Fig. 2A, Supplementary Fig. 3A,D,F). SCALEX successfully preserved the latent structure of the two modalities, but it failed to mix them well (Supplementary Fig. 3C). LIGER and MultiMAP performed the worst in this case, showing total misalignment and chaos in UMAP visualization (Supplementary Fig. 3B,E). Compared to uniPort_cm, Seurat and Harmony, uniPort, which leveraged common, RNA-specific and ATAC-specific genes for integration, demonstrated better separation between CD4 and CD8 cells (Fig. 2A). In order to intuitively show the importance of specific genes, we visualized the gene expression of RNA-specific marker genes INPP4 and FIHT and ATAC-specific marker gene KLRC4.KLRK1 (Fig. 2D,E). These marker genes showed high expression values in CD4 or CD8 cells, which assisted the identification and separation of corresponding cells. From another perspective, the accessibility of DNA to regulatory proteins attaches an essential characteristic which impacts the fate of transcription. We found that the KLRC4.KLRK1 gene has higher DNA accessibility contrast in CD8 Naive T cells (Fig. 2F). This is crucial for cell-type or state identification during integration. We also visualized the confusion matrices between ground truth cell types of scATAC data and predicted annotations through label transfer by a kNN classifier trained with scRNA cell types in the latent space. We compared Seurat, Harmony and SCALEX, but ignored MultiMAP and LIGER owing to their chaotic UMAP visualization, as suggested above (Fig. 2G). Result showed that uniPort identified CD4 Tmem, CD8 Naive and CD Tmem cells better than the other methods, reflecting the contribution of ATAC-specific and RNA-specific genes.

### uniPort integrates scRNA and spatially resolved MERFISH data

We then applied uniPort to integrate scRNA data with high-plex RNA imaging-based spatially resolved MERFISH data, including 30,370 cells with 21,030 genes from six mice measured with dissociated scRNAseq (10X) and 64,373 cells with 155 genes from mouse 1 measured with MERFISH [26]. Concretely, we filtered 153 common genes in both datasets and selected the top 2,000 highly variable genes in scRNA data. It should be noted that the 2,000 genes in scRNA data contain some of the 153 common genes, which are also highly variable genes in scRNA data. Afterwards, we projected cells with common genes of both MERFISH and scRNA data into cell-embedding latent space by uniPort and then reconstructed input and 2000 genes in scRNA from the cell embeddings.

We applied UMAP to visualize the integration results of cell embeddings by uniPort, Harmony, Seurat and SCALEX (Fig. 3A). As shown in the picture, uniPort did a better job of identifying and integrating OD Immature cells, as well as separating them from other cells, with the assistance of 2000 scRNA genes. Both uniPort and SCALEX successfully avoided over-correction by correctly maintaining Ependymal cells apart that were only observed in MERFISH data, demonstrating that uniPort can successfully integrate partially overlapping datasets. We again benchmarked integration performance by Silhouette and Batch Entropy scores, including uniPort, uniPort_cm, SCALEX, Seurat, Harmony and LIGER (Fig. 3B). uniPort, again, outperformed other methods with the highest Silhouette score of 0.669 and the second highest Batch Entropy score of 0.474, slightly below SCALEX of 0.499. uniPort_cm achieved performance very similar to that of uniPort (Silhouette score of 0.665, Batch Entropy score of 0.475). Among all methods, LIGER had the lowest Silhouette score of 0.495, and Seurat had the lowest Batch Entropy score of 0.265. We also computed the ARI, NMI and F1 scores to evaluate clustering performance in latent space (Fig. 3C). Overall, uniPort achieved the highest NMI (0.677), while uniPort_cm ranked first in F1 score (0.786), and SCALEX outperformed other methods in F1 score (0.618).

**Figure 3:**
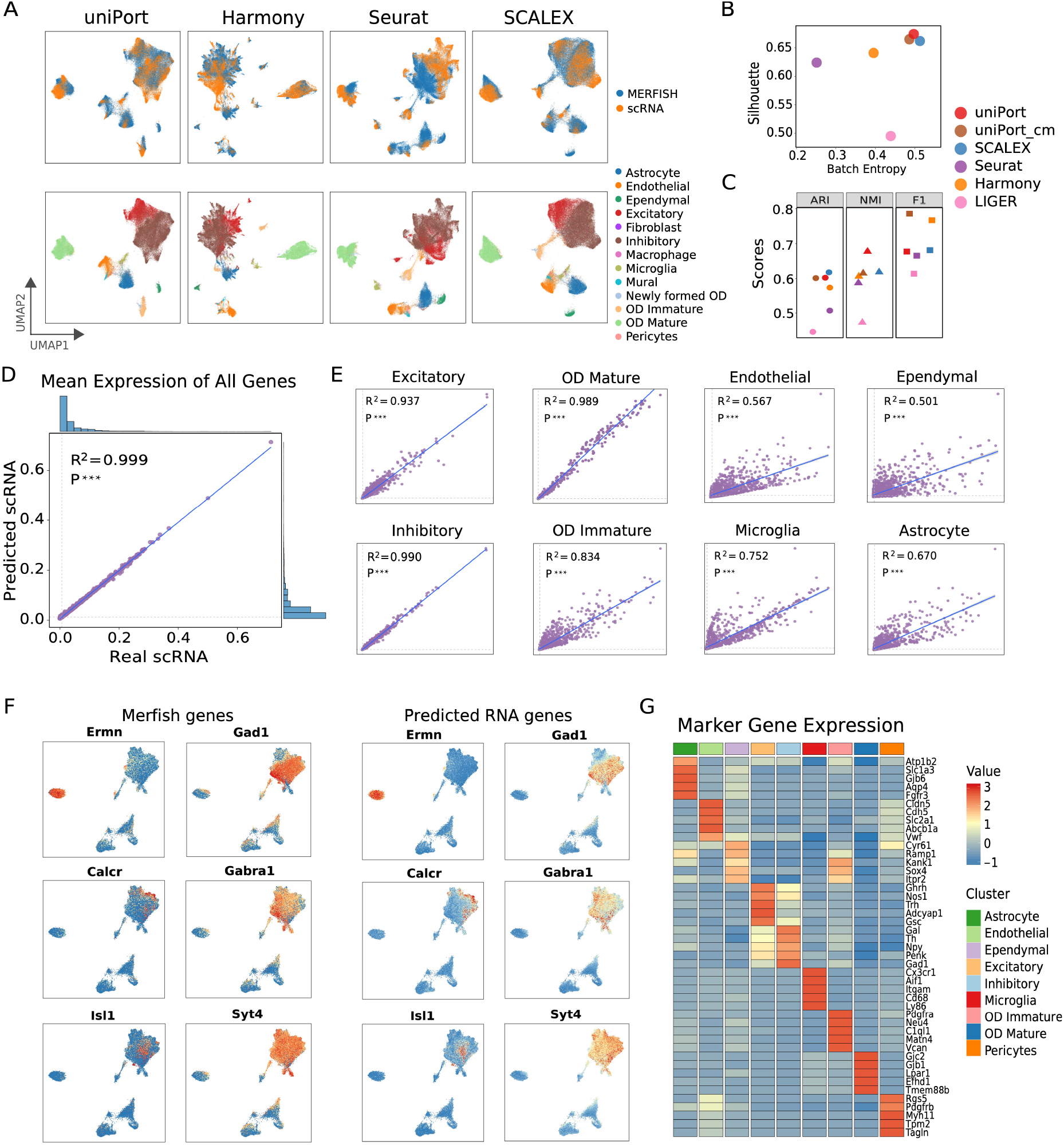
uniPort predicts scRNA genes by MERFISH data. **(A)** UMAP visualization of integration of MERFISH and scRNA data of mouse 1 by uniPort, Harmony, Seurat and SCALEX. **(B)** Comparison of Batch Entropy and Silhouette scores for evaluating integration performance. **(C)** Comparison of ARI, NMI and F1 scores for evaluating clustering performance. **(D)** Correlation of generated and real expression of RNA genes in all cells and in different cell types. **(E)** UMAP visualization of common gene expression in MERFISH and generated RNA data. **(F)** Averaged predicted RNA marker gene expression values through MERFISH data of mouse 2.

### uniPort online predicts genes in scRNA data by a well-trained reference atlas

uniPort trained an encoder network to project cells with common genes across datasets into a common cell-embedding latent space and a decoder network to reconstruct cells with common and specific genes. Therefore, once the coupled-VAE is trained well, it can be regarded as a reference atlas, in turn allowing uniPort to integrate new single-cell data in an online manner without modal retraining. Most importantly, uniPort can generate both common and specific genes in one dataset through common genes in another dataset by the atlas (Supplementary Fig. 4).

To explore uniPort’s ability for online prediction, we utilized the well-trained modal of scRNA and MERFISH data profiled from mouse 1 as a reference atlas and downloaded MERFISH data profiled from mouse 2 with 59,651 cells and 155 genes. We selected the same 153 common genes as in the above-referenced MERFISH data of mouse 2 and input them into the atlas. Then the encoder projected MERFISH data of mouse 2 into the integrated latent space, and the decoder reconstructed the corresponding 2,000 highly variable genes in scRNA from the cell embeddings (Supplementary Fig. 4B). Accordingly, the modal proved to be especially powerful in that it could predict the expression of genes in scRNA data, even through not measured in MERFISH data, in an online manner. The predicted data can be used to enhance the spatial transcriptomics [36, 37].

To assess the validity and qualification of online predicted scRNA data, we calculated the correlation of mean expression of genes in scRNA data and predicted data (Fig. 3D), as well as those genes in different cell types (Fig. 3E), noting that the cell types of predicted scRNA are the same as those in the corresponding MERFISH data. As a result, real and predicted scRNA data were significantly correlated, according to the Pearson correlation coefficient *R*^2^ = 0.999 for all data, and ranged from 0.501 for Ependymal cells to 0.990 for Inhibitory cells. In addition, we plotted marker gene expression values in the UMAP visualization of online projection of MERFISH data of mouse 2 and the same genes in predicted scRNA data, including Ermn, Gad1, Calcr and so on (Fig. 3F). Results showed that the high-expression regions of common genes in MERFISH and generated scRNA were consistent in the latent space. Finally, for verification of cell-type labels, we plotted the top differential marker gene expression for each cell type in predicted scRNA data and observed consistent patterns of cell-type-specific expression (Fig. 3G). For example, Excitatory and Inhibitory cells are difficult to distinguish, since, in essence, they belong to neural subtypes. However, our generated differential marker gene expression exhibited significant differences between the two cell types, such as marker genes Ghrh and Trh in Excitatory cells and Gal and Th in Inhibitory cells.

### uniPort deciphers canonical structures of mouse brain

We next considered the deconvolution of heterogeneous barcoding-based ST data through transferring labels from scRNA data. To estimate cell-type composition for each captured spot and decipher typical organizational structures, we applied uniPort to integrate the anterior slice of adult mouse brain ST data (10X Visium) with 48,721 genes in 2,696 spots and matched brain scRNA data from SPOTlight [28]. uniPort provided a global OT plan, which represents a cell-to-spot probabilistic correspondence between scRNA and ST data, allowing us to deconvolute the proportion of single-cell clusters for ST data according to cell annotations in scRNA data (Supplementary Note 2).

As clear-cut boundaries exhibited, uniPort accurately reconstructed the well-structured layers and deconvoluted 28 cell types (Fig. 4A). Proportion and position of representative clusters, e.g., multiple cortical layers and region-specific cell types, are highly consistent with those of previous studies [28,38]. Despite the complexity of its anatomy, uniPort accurately remodeled and arranged the L2/3-L6 subclusters extending from the boundary to central area (Fig. 4B). In addition, subpopulations of the L6 layer were separated with clear limits, revealing the sensitivity of our method to near-imperceptible signals. The non-neuronal neuroglia cells that provide neurons with support and protection, including astrocytes and oligodendrocytes [39], had corresponding sites paired with their marker genes (e.g., Olig1 and Olig2 of cluster Oligo; Atp1b2 and Slc1a2 of cluster Astro; Dcn and Osr1 of cluster VLMC) (Supplementary Fig. 5). Moreover, vascular and leptomeningeal cells (VLMCs) that form protective sections around the pia membranes of the brain also lay in the border of slice in harmony with anatomical structures [40]. Therefore, our mapping is robust, as demonstrated by either expression of marker genes or anatomy of brain, and can establish an agreement between gene expression-based clustering and anatomical annotation, providing a more thorough and comprehensive understanding than can be achieved through visual inspection.

**Figure 4:**
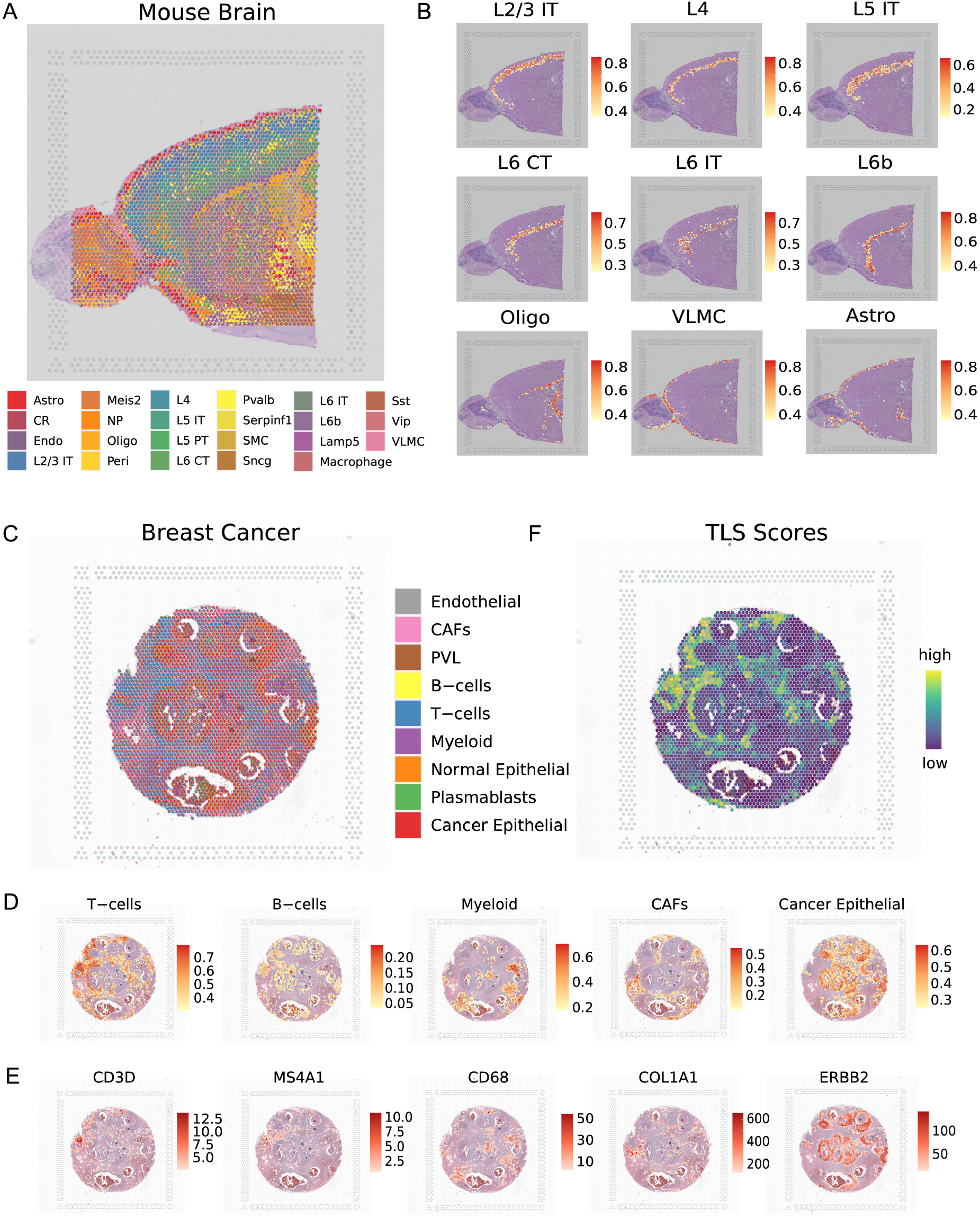
uniPort identifies iconic structure in spatial transcriptomics data (10X Visium). **(A)** Results of mapping spatial data to single-cell data using optimal transport plan. Spatial scatter pie plot displays the well-structured cluster composition in adult mouse brain anterior slice. **(B)** Lists of canonical cerebral cortical neuron type with scaled proportion. **(C)** Spatial deconvolution result of HER2-positive breast cancer. **(D)** Proportion of typical clusters in tumor microenvironment. **(E)** Expression of marker genes corresponding to clusters in (D). **(F)** Tertiary lymphoid structure (TLS) scores inferred from summing the proportion of T cells and B cells together with their colocalization.

### uniPort assists in locating TLS in breast cancer region

The genesis and progression of cancer are generally influenced by their association with the heterogeneous tumor microenvironment (TME) [41] for which ST can provide perfect insight. To further demonstrate its flexible utility, we used uniPort to deconvolute spatial data of HER2-positive breast cancer, containing diffusely infiltrating cells that make it more difficult to deconvolute spots. In total, 2,518 spots with 17,943 genes and 100,064 cells with 29,733 genes [29] were used for integration. As shown in Figure 4, nine main clusters were assigned on spatial images, primarily involving T cells and cancer epithelial cells (Fig. 4C). Moreover, we found that representative clusters scattered in their centralized enrichment region coincided with the area of their marker genes’ expression (Fig. 4D). For example, T and B cells, which establish crucial adaptive immunity through protective immunological memory, were matched well with canonical marker genes, such as CD3D and MS4A1 (Fig. 4D). Myeloid cells, as an innate part of the immune system, also displayed a distribution concordant with the expression of CD68 [42]. Furthermore, cancer epithelial cells protruded along the invasive ductal carcinoma region, corresponding with the expression of ERBB2 as well. Overall, the above results reach concordance between pathological annotation and data-motivated labeling.

Massive research has demonstrated that an increased infiltration of immune cells is highly related to favorable breast cancer prognosis. Tertiary lymphoid structures (TLSs), a kind of ectopic lymph-like organ recently discovered at sites of tumor or inflammation, are considered as a prognostic and predictive factor for cancer patients. Although TLS is inhabited by multiple cell types, its major residents are T and B cells, implicating the TME by their joint colocalization [43]. Through decomposing cell-type proportion of each spot, we can identify TLS signals via colocalization of T and B cells, rendering an identical expression intensity with T and B cells. In general, our approach can harmonize diverse modalities and cater to both high-resolution mapping and recognition of representative architectures across tissues and diseases.

### uniPort reveals cancer heterogeneity in microarray-based spatial data

The area of Visium-based ST data is limited to a 55 *μ*m diameter for each captured spot, which reaches a moderate resolution that translate to 3–30 cells [27]. Latent integration challenges may arise, along with the decrease of spot resolution, as increased mix of ingredients brings more noise. To examine the performance of uniPort in this case, we employed microarray-based ST data of pancreatic ductal adenocarcinoma (PDAC) tissues for integration, the diameter of which stretches for 100 *μ*m [30]. Cell-type deconvolution was applied on 428 spots paired with 1926 single cells, measuring 19,736 genes respectively.

We decomposed 15 main clusters, which exhibit a discrete enrichment and complexity of both normal and tumor composition (Fig. 5A). In detail, normal pancreatic cell types were classified as ductal and acinar cells consistent with the results of previous studies [44], preserving dramatically different distributions and genetic characteristics against those of cancer cells. As for malignant pancreatic cells, we grouped as cancer clone A and B clusters on the basis of genetic differences. Once again, histological annotations of normal and cancerous regions were, overall, in line with their data-driven labels (Fig. 5B), and essential constituents of the TME were indicated by their marker genes (Fig 5C,D).

**Figure 5:**
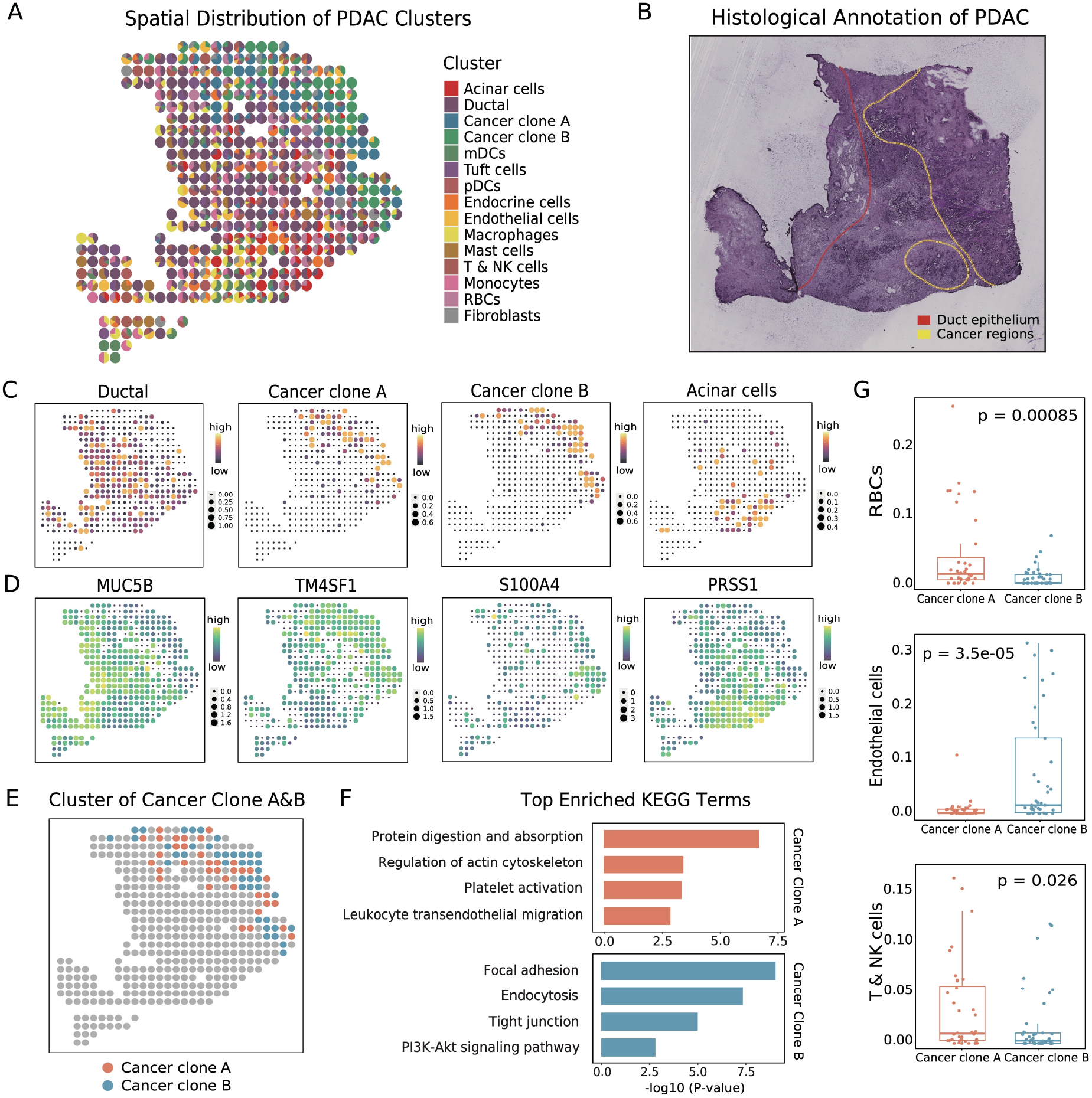
uniPort identifies distinct cancer subtypes in microarray-based spatial data. **(A)** Spatial deconvolution results of pancreatic ductal adenocarcinoma (PDAC). **(B)** Annotation of PDAC tumor cryosection on ST slide. The red line circles ductal epithelium region, and the yellow line circles the cancer region. **(C)** Proportion and distribution of typical clusters in PDAC. **(D)** Expression of marker genes corresponding to clusters in (C). **(E)** Distribution of cancer clone subtypes. **(F)** Top enriched KEGG terms of distinct cancer subtypes. **(G)** Significant differences of cluster composition between the two cancer clone regions.

To gain further insight into the heterogeneity of cancer subtypes, we confirmed their identity, accounting for maximum proportion of each spot (Fig. 5E). Top enrichment KEGG pathways isolated them into distinct functional assemblies (Fig. 5F). Cancer clone A is suspected to be an invasive phenotype attributed to high enrichment of platelet activation and leukocyte transendothelial migration pathway. Lumps of data have proved that platelets are closely related to high risk of metastasis in patients with pancreatic cancer [45]. Furthermore, the proportion of hematogenous cells, including red blood cells (RBCs), T cells and natural killer (NK) cells, in cancer clone A significantly increased (Fig. 5G), which is consistent with the results of functional analysis. Tight junction plays a critical regulatory role in physiologic secretion of pancreas, and its disruption contributes to the pathogenesis of progressive pancreatic cancer [46]. Furthermore, PI3K signaling can potentially modify the TME to dictate outcome, which must be considered in order to have therapeutic opportunities for targeting PDAC [47]. All these beneficial signals were enriched in the cancer clone B region where endothelial cells showed a significant presence, suggesting a less malignant cluster in contrast with cancer clone A. In sum, then, our method can manipulate an extensive application spectrum of varying resolutions, revealing the subtle heterogeneous TME.

## Discussion

We introduce a unified deep learning method named uniPort for single-cell data integration and apply it to integrate transcriptomic, epigenomic, spatially resolved high-plex RNA imaging- and barcoding-based single-cell data. uniPort combines a coupled-VAE and Minibatch-UOT and leverages both highly variable common and dataset-specific genes for integration. It is a nonlinear method that projects all datasets into a common latent space and outputs their latent representations between datasets, enabling both visualization and downstream analysis.

Generally, uniPort tackles several computational challenges, starting with removing the constraint of paired cells found in other autoencoder-based modals by employment of Minibatch-UOT. Different from existing methods that only consider common genes across datasets, we also take advantage of genes unique to each dataset, typically capturing cell-type heterogeneity not present in the common genes, especially when the number of common genes is small. Besides, uniPort shows its power and potential for online prediction of unmeasured modalities through constructing a reference atlas owing to the generalization ability of coupled-VAE. It is relevant to point out that uniPort can even predict unique genes in one dataset through common genes in another dataset without having to train from scratch. Moreover, uniPort can optionally output a global OT plan for downstream analysis, such as flexible label transfer learning, for deconvolution of spatial heterogeneous data.

Although many mechanisms are involved, uniPort is still computationally efficient and scalable to integrate large-scale and partially overlapping datasets, which is difficult for other OT-based methods. We demonstrate that uniPort achieves the highest values in tested scores, provides better joint visualization than other methods, and successfully deconvolutes spatial heterogeneous data using OT space. Finally, with the rapid development of paired-cell datasets and various heterogeneous modalities, we also demonstrate the generalizability of uniPort to other types of single-cell data by integrating datasets profiled from the same cells or datasets without aligned common genes (Supplementary Notes 3 and 4, Supplementary Figs. 6 and 7). With no inherent reliance on any prior information, our framework offers the flexibility necessary to match prior information, e.g., cell-type annotations or cell-cell correspondence, when available (Supplementary Note 5). We will keep updating and improving the framework in anticipation that uniPort will find a wide range of applications in the area of integrative multi-omics data analysis.

## Methods

### uniPort framework

uniPort inputs each dataset though a coupled variational autoencoder (coupled-VAE) framework and learns its *K*-dimensional features. Given a sample ***x***, the corresponding *K*-dimensional latent vector ***z*** can be obtained by a variational posterior *p*(***z***|***x***) approximated by a probabilistic encoder *ψ*(***z***|***x***). Conversely, a probabilistic decoder *ϕ*(***x***|***z***) produces a distribution over the possible corresponding values of ***x*** given ***z***. The coupled-VAE jointly learns *ψ* and *ϕ* to maximize the evidence lower bound (ELBO) with a balanced parameter *λ*:

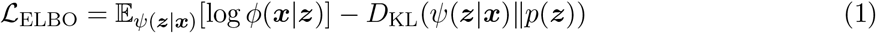

The ELBO consists of a reconstruction term which encourages the output data to be similar to the input data, in addition to a Kullback-Leibeler divergence regularization term which regularizes the variational posterior to follow the prior distribution *p*(***z***). We set the prior to be the centered isotropic multivariate Gaussian 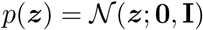 and the variational posterior to be a multivariate Gaussian with a diagonal covariance structure 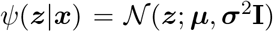, where the mean vector ***μ*** and standard deviation vector ***σ*** are outputs of the encoder. Then, the latent vector ***z*** can be obtained through reparameterization by ***z*** = ***μ*** + ***σ*** × *ϵ*, where *ϵ* is sampled from 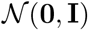.

Here, we formulate our method for the case of two datasets, while it can be easily generated for cases of multiple single-cell datasets. Suppose there are two single-cell datasets **X** = {***x***_1_,…, ***x**_n_x__*} with *n_x_* cells and *d_x_* genes and **Y** = {***y***_1_,…, ***y**_n_y__*} with *n_y_* cells and *d_y_* genes. We filter out the top *k* highly variable common genes in both datasets as **X***^c^* and **Y***^c^* and the top *k_x_* and *k_y_* highly variable genes in **X** and **Y** as **X***^s^* and **Y***^s^*, individually. We project both **X***^c^* and **Y***^c^* into a generalized cell-embedding latent space using a dataset-free probabilistic encoder *ψ* and a decoder *ϕ* with Dataset-Specific Batch Normalization (DSBN) layers [23]. Then, to leverage the information of dataset-specific highly variable genes, we also introduce two decoders, *ϕ_x_* and *ϕ_y_*, to reconstruct **X***^s^* and **Y***^s^* from the latent variables, as well. Overall, the modified ELBO* loss for coupled-VAE is given by

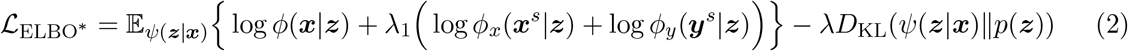

where ***x*** ∈ **X***^c^* ∪ **Y***^c^*, ***x**^s^* ∈ **X***^s^*, ***y**^s^* ∈ **Y***^s^*, and *λ*_1_, *λ*_2_ are balanced parameters. We set *λ*_1_ = 0.5 for all experiments in this paper.

To better integrate heterogeneous single-cell datasets in the latent space, we design an alignment term for different datasets using Minibatch Unbalanced Optimal Transport (Minibatch-UOT) [22]. uniPort computes the Minibatch-UOT loss between datasets **X** and **Y** described as follows. For two K-dimensional Gaussian distributions 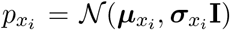 and 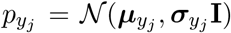 corresponding to cell ***x**_i_* and ***y**_j_*, where ***μ**_x_i__*, 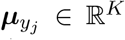 and ***σ**_x_i__*, 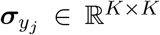 represent the output mean and covariance vectors by encoder *ψ*, respectively. The Minibatch-UOT cost between cell ***x**_i_* and ***y**_j_* is defined as (Supplementary Note 1):

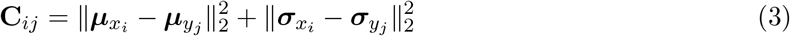

We then compute the following optimal Minibatch-UOT plan [22] with batch size *B_x_* and *B_y_*:

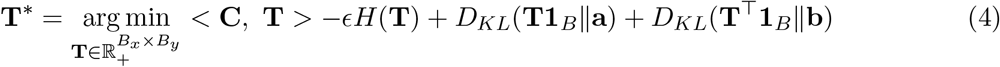

where < **C**, **T** >= ∑*_ij_* **C***_ij_* **T***_ij_*, entropy regularization term *H*(**T**) = – ∑*_ij_* **T***_ij_*(log **T***_ij_* – 1), and 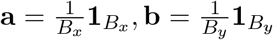. *ϵ* is a balanced parameter. Equation (4) is a strictly convex optimization problem and can be solved via an efficient inexact proximal point method (IPOT) [48] as

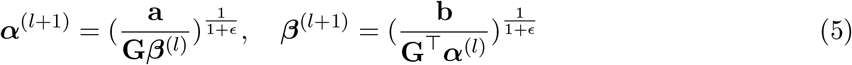

starting from 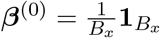, where 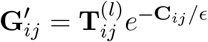 and the optimal Minibatch-UOT plan 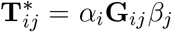. Therefore, the Minibatch-UOT loss is given by

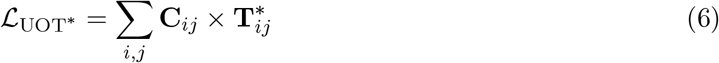

The total loss function minimized by uniPort is formulated as

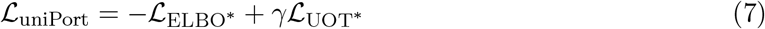

Once **T**\* is obtained, we provide users an option to output a global OT plan **T***_global_* for tasks which need cell-cell probabilistic correspondence. Specifically, we first initialize 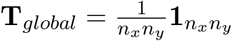. After each calculation of optimal Minibatch-UOT plan **T**\*, we update **T***_global_* by replacing rows and columns by **T**\* which is sampled by the mini-batch strategy. Note that storing a global dense matrix **T***_global_* needs more memory. Therefore, computing capacity with respect to memory should be ascertained before using this option. We also provide an option for user-guided sample weights if cells are not uniformly matched. In this case, we set vectors **a** and **b** in Minibatch-UOT as weighted vectors specified by users, instead of uniform vectors, and reweight reconstruction loss in coupled-VAE, as well (Supplementary Note 6).

### uniPort algorithm

To integrate two single-cell datasets **X** = {***x***_1_,…, ***x**_n_x__*} and **Y** = {***y***_1_,…, ***y**_n_y__*}, uniPort performs the following steps:

1. Select the top *k* highly variable common genes in **X** and **Y**, formulate as **X***^c^* and **Y***^c^*, and top *k_x_* and *k_y_* highly variable genes (may also contain some common genes) in **X** and **Y**, individually, and then formulate as **X***^s^* and **Y***^s^*.
2. Initialize coupled-VAE encoder *ψ* and decoders *ϕ, ϕ_x_, ϕ_y_*, and a global OT plan 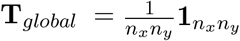 (optional, depending on computing memory).
3. For *m* ← 1,…, *M* do the following

a. Randomly sample an integer index set 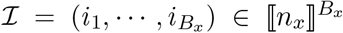 for dataset **X**, and 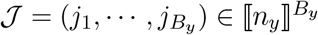 for dataset **Y** without replacement, respectively.
b. Initialize Minibatch-UOT plan 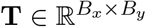 by sampling rows and columns of **T***_global_*, corresponding to 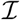 and 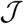, or uniform distribution when **T***_global_* is not specified.
c. Input both 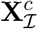 and 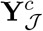 through the shared probabilistic encoder *ψ* to obtain (***μ**_x_, **σ**_x_*) and (***μ**_y_, **σ**_y_*) and then reparameter ***z**_x_* and ***z**_y_* by (***μ**_x_, **σ**_x_*) and (***μ**_y_, **σ**_y_*);
d. Reconstruct 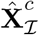 and 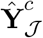 by decoder *ϕ*, and 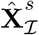 and 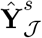 by decoders *ϕ_x_* and *ϕ_y_*, from ***z**_x_* and ***z**_y_*.
e. Compute mini-batch transport cost **C** via (3) and obtain optimal Minibatch-UOT plan **T**\* via (4).
f. Fix **T**\* and update *ψ, ϕ, ϕ_x_* and *ϕ_y_* through back propagation via minimizing 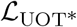 and 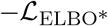.
g. Update **T***_global_* with rows and columns as **T**, corresponding to 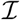 and 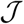 (optional).

### Training details

uniPort consists of one encoder and three decoders for integration of two datasets. The encoder is a two-layer neural network (1024-128) with the ReLU activation function. The decoders have no hidden layer, but directly connect the 16-dimensional latent variables to the output layers with the Sigmoid activation function. The Adam optimizer with a 5e-4 weight decay is used to maximize the ELBO. Mini-batch size is 256. We set all the training with learning rate to 2e-4 and optimal transport entropy regularization parameter *ϵ* to 0.1. We choose balance parameters for training coupled-VAE and Minibatch-UOT from *λ* ∈ {0.5, 1.0, 5.0}, *γ* ∈ {0.5, 1.0} for all datasets. uniPort is robust to different choices of *λ* and *γ* within a certain range and scalable to large-scale data (Supplementary Note 7 and Supplementary Fig. 8). The default maximum number of training iterations is 60,000, and an early stopping is triggered when no improvement has occurred for 10 epochs. Our experimental environment includes an AMD EPYC 7302 16-Core Processor, 256GB DDR4 memory and NVIDIA GPU Tesla T4. See Supplementary Note 8 for details of other methods.

### Data preprocessing

We preprocessed data as follows: (1) We filtered out cells with fewer than 200 genes and filtered out genes observed in fewer than 3 cells for PBMC data. No cells or genes were filter out for other data. (2) We normalized total counts of each cell to 10,000. (3) We performed log-normalization of all datasets with an offset of 1. (4) We identified *k* = 2, 000 highly variable common genes across cells of all datasets. We also identified *k_x_* = *k_y_* = 2, 000 highly variable genes for each dataset, respectively. (5) We normalized values of each gene to the range of 0-1 within each dataset by the *MaxAbsScaler* function in the *scikit-learn* package in Python. The processed matrix was used as input for uniPort. For scATAC-seq data, a gene activity matrix was created by the *GeneActivity* function in the Signac R package to quantify the activity of each gene from scATAC-seq data. The subsequent preprocessing of gene activity matrix followed the same preprocessing as above.

## Data availability

All data analyzed in this article are publicly available through online sources. We present links to all data sources in Supplementary Table 1.

## Code availability

uniPort software is available at https://github.com/caokai1073/uniPort.

## Acknowledgements

This work was supported by the National Key R&D Program of China under Grant 2019YFA0709501, NSFC grants (Nos. 61733018, 12071466, 62173250), Shanghai Municipal Science and Technology Major Project (No. 2021SHZDZX0100), the Fundamental Research Funds for the Central Universities, and LSC of CAS. Fig. 1A was created with BioRender.com.

