## Supplementary Notes and Figures for "uniPort: a unified computational framework for single-cell data integration with optimal transport"

#### 1. Mini-batch unbalanced optimal transport

We first give a brief introduction to optimal transport (OT), which has attracted more and more attention in analysis of single-cell datasets recently. Suppose we have two single-cell datasets $\mathbf{X} = \{\boldsymbol{x}_i\}_{i=1}^{n_x}, \boldsymbol{x}_i \in \mathbb{R}^K$ and $\mathbf{Y} = \{\boldsymbol{y}_j\}_{j=1}^{n_y}, \boldsymbol{y}_j \in \mathbb{R}^K$ in the latent $K$-dimensional spaces. Two discrete measures with weights $\mathbf{a}$, $\mathbf{b}$ are defined as

$$\mathbf{a} = \sum_{i=1}^{n_x} a_i \delta_{\boldsymbol{x}_i}, \ \mathbf{b} = \sum_{i=1}^{n_y} b_i \delta_{\boldsymbol{y}_i} \tag{1}$$

where $\delta_{\boldsymbol{x}}$ is the Dirac at cell $\boldsymbol{x}$, and $\mathbf{a}$, $\mathbf{b}$ belong to the probability simplex: $\Sigma_n \stackrel{\text{def}}{=} \{\mathbf{a} \in \mathbb{R}_+^n : \sum_{i=1}^n \mathbf{a}_i = 1\}$. Typically, we initialize $\mathbf{a}$, $\mathbf{b}$ as uniform distribution with each element be the same, that is $\mathbf{a} = \frac{1}{n_x}\mathbf{1}_{n_x}, \mathbf{b} = \frac{1}{n_y}\mathbf{1}_{n_y}$. However, we also provides a user-guided reweight option in our method if the importance of each cell for alignment is not the same (Supplementary Note 6).

Besides, the OT plan, i.e., cell-cell probabilistic coupling matrix, $\mathbf{T} \in \mathbb{R}_+^{n_x \times n_y}$, as well as a cost matrix $\mathbf{C} \in \mathbb{R}_+^{n_x \times n_y}$, describes the probabilities of aligning cells and the distances between cells across datasets in the latent space, respectively. Consequently, the discrete OT problem is defined as follows

$$\boldsymbol{L}(\mathbf{a}, \mathbf{b}) \stackrel{\text{def}}{=} \min_{\mathbf{T} \in \Pi(\mathbf{a}, \mathbf{b})} \langle \mathbf{C}, \mathbf{T} \rangle = \min_{\mathbf{T} \in \Pi(\mathbf{a}, \mathbf{b})} \sum_{i,j} \mathbf{C}_{ij} \mathbf{T}_{ij} \tag{2}$$

where $\Pi(\mathbf{a}, \mathbf{b}) \stackrel{\text{def}}{=} \{\mathbf{T} \in \mathbb{R}_+^{n_x \times n_y} : \mathbf{T}\mathbf{1}_{n_y} = \mathbf{a}, \ \mathbf{T}^\top \mathbf{1}_{n_x} = \mathbf{b}\}$, and $\langle \cdot, \cdot \rangle$ denotes Frobenius dot product. Besides, in order to introduce some smoothness to the transport matrix, the approximate solution according to the regularization parameter $\epsilon$ writes as

$$\boldsymbol{L}_\epsilon(\mathbf{a}, \mathbf{b}) \stackrel{\text{def}}{=} \min_{\mathbf{T} \in \Pi(\boldsymbol{a}, \boldsymbol{b})} \langle \mathbf{C}, \mathbf{T} \rangle - \epsilon \sum_{i,j} \mathbf{T}_{i,j} \log(\mathbf{T}_{i,j}) \tag{3}$$

Equation (3) is strictly convex optimization problem and can be solved efficiently via iterative Bergman projections [1]:

$$\boldsymbol{\alpha}^{(l+1)} = \frac{\mathbf{a}}{\mathbf{G}\boldsymbol{\beta}^{(l)}}, \quad \boldsymbol{\beta}^{(l+1)} = \frac{\mathbf{b}}{\mathbf{G}^\top \boldsymbol{\alpha}^{(l)}} \tag{4}$$

starting from $\boldsymbol{\beta}^{(0)} = \frac{1}{n_y}\mathbf{1}_{n_y}$, where $\mathbf{G}_{ij} = e^{-\mathbf{C}_{ij}/\epsilon}$, and the optimal transport plan $\mathbf{T}^*_{ij} = \alpha_i \mathbf{G}_{ij}\beta_j$. In uniPort, we employ a more robust and efficient inexact proximal point method (IPOT) [7] to compute the OT plan. Concretely, uniPort replaces $\mathbf{G}_{ij}$ with $\mathbf{G}'_{ij} = \mathbf{T}^{(l)}_{ij} e^{-\mathbf{C}_{ij}/\epsilon}$ in Eq. (4).

To combine OT with coupled-VAE, we utilizes the mini-batch unbalanced optimal transport (Minibatch-UOT) [4], which is a geometrically robust version. Minibatch-UOT is computed between mini batches which is practical for large-scale datasets and deep learning applications, and also decreases the influence of undesired outliers which makes it suitable for partially-overlap datasets. Compared to classic OT, Minibatch-UOT changes the Eq. (3) as

$$\min_{\mathbf{T}\in\mathbb{R}_+^{B_x\times B_y}} < \mathbf{C},\ \mathbf{T} > -\epsilon H(\mathbf{T}) + \tau \left( D_{KL}(\mathbf{T}\mathbf{1}_B \| \mathbf{a}) + D_{KL}((\mathbf{T})^\top \mathbf{1}_B \| \mathbf{b}) \right) \tag{5}$$

where $B_x$ and $B_y$ are mini-batch sizes of datasets $\mathbf{X}$ and $\mathbf{Y}$, and $\mathbf{T}$ and $\mathbf{C}$ are mini-batch OT plan and cost. $D_{KL}$ is KL divergence, and $\tau$ is a marginal penalization. It should be noted that when $\tau \to \infty$, the algorithm degenerates into balanced OT. We set $\tau = 1$ in all experiments. Therefore, the corresponding Eq. 4 can be rewritten as

$$\boldsymbol{\alpha}^{(l+1)} = \left(\frac{\mathbf{a}}{\mathbf{G}\boldsymbol{\beta}^{(l)}}\right)^{\frac{1}{1+\epsilon}}, \quad \boldsymbol{\beta}^{(l+1)} = \left(\frac{\mathbf{b}}{\mathbf{G}^\top \boldsymbol{\alpha}^{(l)}}\right)^{\frac{1}{1+\epsilon}} \tag{6}$$

Besides, uniPort computes the OT cost $\mathbf{C}$ between the Gaussian mixture models $\{\mathcal{N}(\boldsymbol{\mu}_{x_k}, \boldsymbol{\sigma}^2_{x_k}\mathbf{I})\}_{k=1}^{B_x}$ and $\{\mathcal{N}(\boldsymbol{\mu}_{y_k}, \boldsymbol{\sigma}^2_{y_k}\mathbf{I})\}_{k=1}^{B_y}$, instead of latent vectors $\boldsymbol{z}_x$ and $\boldsymbol{z}_y$, where $\boldsymbol{\mu}$ and $\boldsymbol{\sigma}$ are output of the probabilistic encoder of coupled-VAE.

**Definition 1** *For two $K$-dimensional Gaussian distributions $\mathcal{N}(\boldsymbol{\mu}_x, \boldsymbol{\Sigma}_x)$ and $N(\boldsymbol{\mu}_y, \boldsymbol{\Sigma}_y)$. The Wasserstein distance admits a closed-form expression [6]:*

$$W(p_x, p_y) = \|\boldsymbol{\mu}_x - \boldsymbol{\mu}_y\|^2 + trace\left(\boldsymbol{\Sigma}_x + \boldsymbol{\Sigma}_y - 2(\boldsymbol{\Sigma}_x^{\frac{1}{2}}\boldsymbol{\Sigma}_y\boldsymbol{\Sigma}_x^{\frac{1}{2}})^{\frac{1}{2}}\right) \tag{7}$$

*When the covariance matrices are diagonal, i.e., $\boldsymbol{\Sigma} = diag(\boldsymbol{\sigma}^2)$, where $\boldsymbol{\sigma} \in \mathbb{R}^K$ is the standard deviation vectors, we can rewritten Wasserstein distance as [8]:*

$$W(p_x, p_y) = \|\boldsymbol{\mu}_x - \boldsymbol{\mu}_y\|^2 + \|\boldsymbol{\sigma}_x - \boldsymbol{\sigma}_y\|^2 \tag{8}$$

Therefore, according to Definition 1, the optimal transport cost $\mathbf{C}_{ij}$ between $i$-th component in $\{\mathcal{N}(\boldsymbol{\mu}_{x_k}, \boldsymbol{\sigma}^2_{x_k}\mathbf{I})\}_{k=1}^{B_x}$ and $j$-th component in $\{\mathcal{N}(\boldsymbol{\mu}_{y_k}, \boldsymbol{\sigma}^2_{y_k}\mathbf{I})\}_{k=1}^{B_y}$ is defined as

$$\mathbf{C}_{ij} = \|\boldsymbol{\mu}_{x_i} - \boldsymbol{\mu}_{y_j}\|_2^2 + \|\boldsymbol{\sigma}_{x_i} - \boldsymbol{\sigma}_{y_j}\|_2^2 \tag{9}$$

We provide a Python package for the implementation of uniPort at `https://github.com/caokai1073/uniPort`. Parts of the code are based on modifications of SCALEX (`https://github.com/jsxlei/SCALEX`) and RAE (`https://github.com/HongtengXu/Relational-AutoEncoders`).

### 2. Global OT plan for high-plex RNA imaging-based and barcoding-based ST data

uniPort can output a global OT plan, i.e., cell-to-spot probabilistic matching matrix, that flexible transfer labels for deconvolution of spatial heterogeneous data, such as high-plex RNA imaging-based and low-resolution microarray-based ST data. After obtaining the OT plan, we need to perform some preprocessing step first. Concretely, we first sum the transport mass, i.e., probability, of every cluster in scRNA data for each spot, which defined as a matrix $\mathbf{M}$. If sample-reweight option in Supplementary Note 6 is chosen, we multiply $\mathbf{M}$ with the cluster proportion of scRNA data. Otherwise, we average $\mathbf{M}$ by divided by number of cells belong to each cluster. For a more flexible application, $\mathbf{M}$ can also be filtered by top percent clusters according to user's requirement.

### 3. uniPort integrates datasets profiled from the same cells

We also develop a method in uniPort for integrative clustering of multiple datasets simultaneously profiled from the same cells, named as uniPort_v (Supplementary Figure 6). uniPort_v aims to improve the clustering performance of one modality with the help of other modalities. It takes one dataset as input and projects the data into a latent space using a encoder. Then, uniPort_v reconstructs different modalities through different decoders, which revises the clustering performance in the latent space. We tested the clustering performance of uniPort_v with CITE and scRNA datasets. Results showed that without the integration of CITE data, the Silhouette score of scRNA data in the latent space through encoder was 0.621 (Supplementary Figure 6B). However, if we involved the CITE data, the Silhouette score increased to 0.680 (Supplementary Figure 6C). We also tested the change of the Silhouette score with different balanced parameter $\lambda_1$, which reflects the extent of importance of CITE data (Supplementary Figure 6D). The curve showed that with the increase of $\lambda_1$, the Silhouette increased to 0.680 ($\lambda_1 = 0.2$) first and then decreased. We suggest the parameter to be set from 0 to 0.5.

### 4. uniPort integrates datasets without aligned common genes

uniPort framework involves a method for integration of datasets without common genes, marked as uniPort_d for convenience (Supplementary Figure 7). uniPort_d takes two single-cell datasets with distinct features as input, which means there is no common gene as reference for alignment. We employ two dataset-specific encoders $\psi_x$ and $\psi_y$ to project the datasets into the same dimensional cell-embedding latent space and perform Minibatch-UOT to align the cells. Afterwards, two dataset-specific decoders $\phi_x$ and $\phi_y$ are used to reconstruct the inputs, respectively.

We tested the performance of uniPort_d with mouse spleen and PBMC datasets without aligning common genes. The results of mouse spleen showed that uniPort_d successfully integrated main cell types, including B cells, Granulocyte, Macrophage and NK, but failed to distinguish T CD4 and T CD8 cells (Supplementary Figure 7C). For PBMC, uniPort_d accurately integrated CD4 Naive, CD 14 cells, but failed on other cell types (Supplementary Figure 7D), which is reasonable owing to the lack of reference information, neither from cells nor genes. The Silhouette and Batch Entropy scores for mouse spleen are 0.508 and 0.462, and for PBMC are 0.618 and 0.613 (Supplementary Figure 7B), which are dramatically lower than uniPort using common genes.

### 5. Contrastive learning with reference guided prior information

To improve the performance of integration, we also develop a contrastive learning [5] method to incorporate cell type annotations or any cell-cell correspondence if available, which was introduced by our former work Pamona [2] and also proposed by MAT$^2$ [9]. Follow the definition in MAT$^2$, for cell $x_i$ in dataset $\mathbf{X}$ and cell $y_j$ in reference dataset $\mathbf{Y}$, if they have the same cell type annotation or a prior correspondence, we regard the cell triplets as a positive anchor, and negative conversely. We combine cell triplets with Minibatch-UOT, which is similar to the Joint Distribution Optimal Transport (JDOT) [3]. Specifically, we define a prior matrix $\mathbf{F} \in \mathbb{R}^{B_x \times B_y}$ where

$$\mathbf{F}_{ij} = \begin{cases} \alpha, & \text{if triplet } (i,j) \text{ is negative} \\ 1, & \text{if triplet } (i,j) \text{ is unknown} \\ 1/\alpha, & \text{if triplet } (i,j) \text{ is positive.} \end{cases} \tag{10}$$

Here $\alpha \geq 1$ is a user-guided parameter reflecting the confidence of cell type annotations or correspondence, and larger $\alpha$ means better confidence. We multiply $\mathbf{F}$ with $\mathbf{C}$ to formulate the final transport cost

$$\mathbf{C}_{ij} \leftarrow \mathbf{C}_{ij} * \mathbf{F}_{ij} \tag{11}$$

We added the contrastive learning to uniPort_d and renamed it as uniPort_d_cl, and apply it to mouse spleen and PBMC datasets as well (Supplementary Fig. 7E, F). Results showed that with the help of guided information of cell-type annotations, uniPort_d_cl dramatically improved the performance of integration. For example, the Silhouette score and Batch Entropy increased to 0.578 and 0.505 for mouse spleen, and 0.750 and 0.583 for PBMC.

### 6. Sample reweight during integration

Most single-cell data integrative methods give cells the same importance during integration. However, in some real-world tasks, rare cells in one modality deserve much attention and should be matched with massive cells in other modalities. Therefore, we provide an option for user-guided sample weights if cells should not be uniformly matched. In this case, we set **a** and **b** in Mini-batch UOT as weighted distributions **p** and **q** specified by users instead of uniform vectors, and reweight reconstruction loss in coupled-VAE as well. Specifically, if users want to give samples in dataset $\mathbf{X}$ a new weight distribution $\mathbf{p}$ and samples in dataset $\mathbf{Y}$ a new $\mathbf{q}$, then the Minibatch-UOT loss becomes

$$\min_{\mathbf{T} \in \mathbb{R}_+^{B_x \times B_y}} < \mathbf{C},\ \mathbf{T} > -\epsilon H(\mathbf{T}) + \tau \left( D_{KL}(\mathbf{T}\mathbf{1}_B \| \mathbf{p}) + D_{KL}((\mathbf{T})^\top \mathbf{1}_B \| \mathbf{q}) \right) \tag{12}$$

Besides, the ELBO loss becomes

$$\mathcal{L}_{\text{ELBO}^*} = \mathbb{E}_{\psi(\boldsymbol{z}|\boldsymbol{x})} \left\{ \frac{\mathbf{r}}{2} \log \phi(\boldsymbol{x}|\boldsymbol{z}) + \lambda_1 \Big( \mathbf{p} \log \phi_x(\boldsymbol{x}^s|\boldsymbol{z}) + \mathbf{q} \log \phi_y(\boldsymbol{y}^s|\boldsymbol{z}) \Big) \right\} - \lambda_2 D_{\text{KL}}(\psi(\boldsymbol{z}|\boldsymbol{x}) \| p(\boldsymbol{z})) \tag{13}$$

where $\boldsymbol{x} \in \mathbf{X}^c \cup \mathbf{Y}^c, \boldsymbol{x}^s \in \mathbf{X}^s$ and $\boldsymbol{y}^s \in \mathbf{Y}^s$, and $r = [\mathbf{p}\ ;\ \mathbf{q}]$ represents concatenation of two weight distributions.

### 7. Computational cost

We tested the scalability of uniPort to large-scale data by measuring both maximum memory usage and total runtime (Supplementary Figure 8). To test uniPort's scalability against other methods, we sampled PBMC data (11,259 cells) to creative seven benchmark datasets with $5K$, $10K$, $20K$, $40K$, $80K$, $160K$ and $320K$ cells. uniPort, SCALEX and Harmony scaled well beyond $320K$ cells, while MultiMAP, Seurat and LIGER is not suitable for integration of datasets beyond $160K$ cells in this case. In sum, the mini-batch strategy in deep learning framework dramatically reduced uniPort's time and memory consumption. For example, uniPort required only 19.4 gigabytes (GB) for $320K$ cells, better than other methods except Harmony. Although uniPort consumed more runtime when cell number is less than $20K$ cells, which was about twice that of SCALEX owing to the addition of optimal transport computation, it was still very efficient and consumed almost constant runtime (about 11 minutes) with the cell number increased.

### 8. Settings used in other methods

We benchmarked uniPort against five other methods, including Seurat v4.1.0, LIGER v1.0.0, Harmony v1.0, MultiMAP v1.0.3 and SCALEX v0.2.0. We follow the suggested integration tutorial of each method, and set all hyperparameters as default.

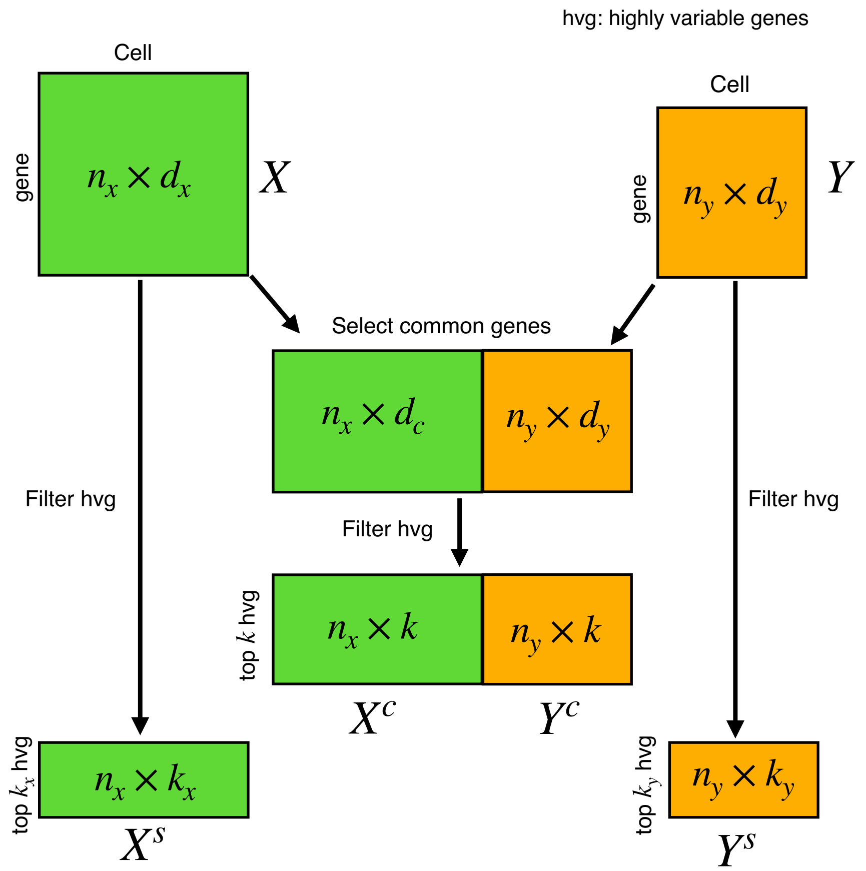


**Supplementary Figure 1. Gene selection.** Steps for selecting $k$ common highly variable genes (hvg), and $k_x$ dataset-specific hvg of dataset $\mathbf{X}$ and $k_y$ dataset-specific hvg of dataset $\mathbf{Y}$.

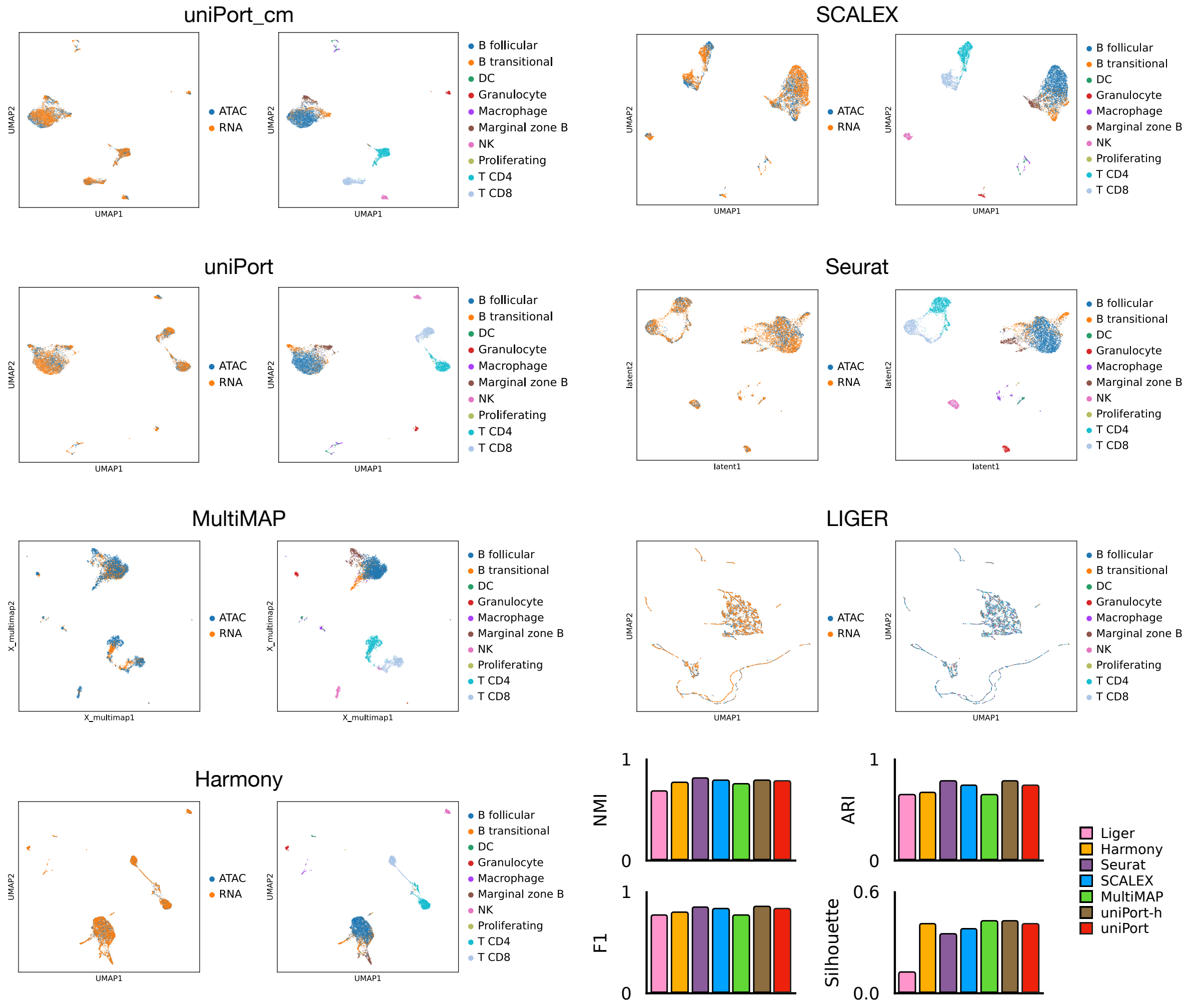


**Supplementary Figure 2. Integration of mouse spleen.** Integration results of mouse spleen datasets by different methods, including uniPort, uniPort_cm, LIGER, SCALEX, Seurat, MultiMAP and Harmony.

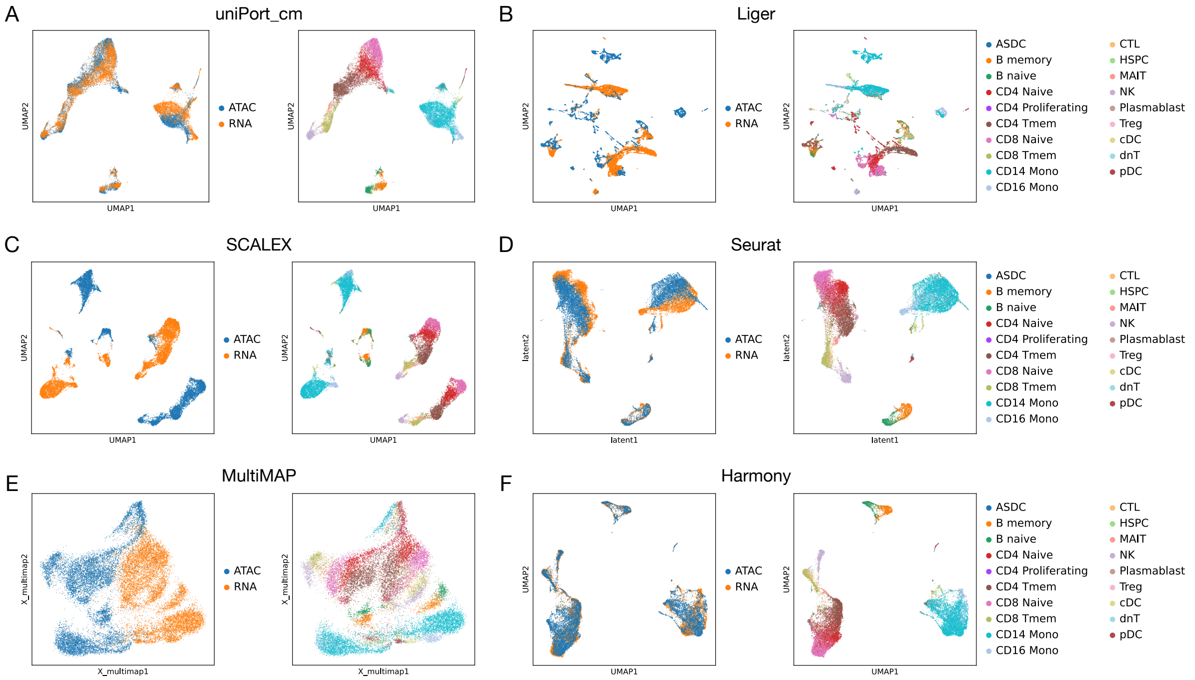


**Supplementary Figure 3. UMAP visualization.** UMAP visualization of integrative results of PBMC data by different methods, including uniPort_cm, LIGER, SCALEX, Seurat, MultiMAP and Harmony.

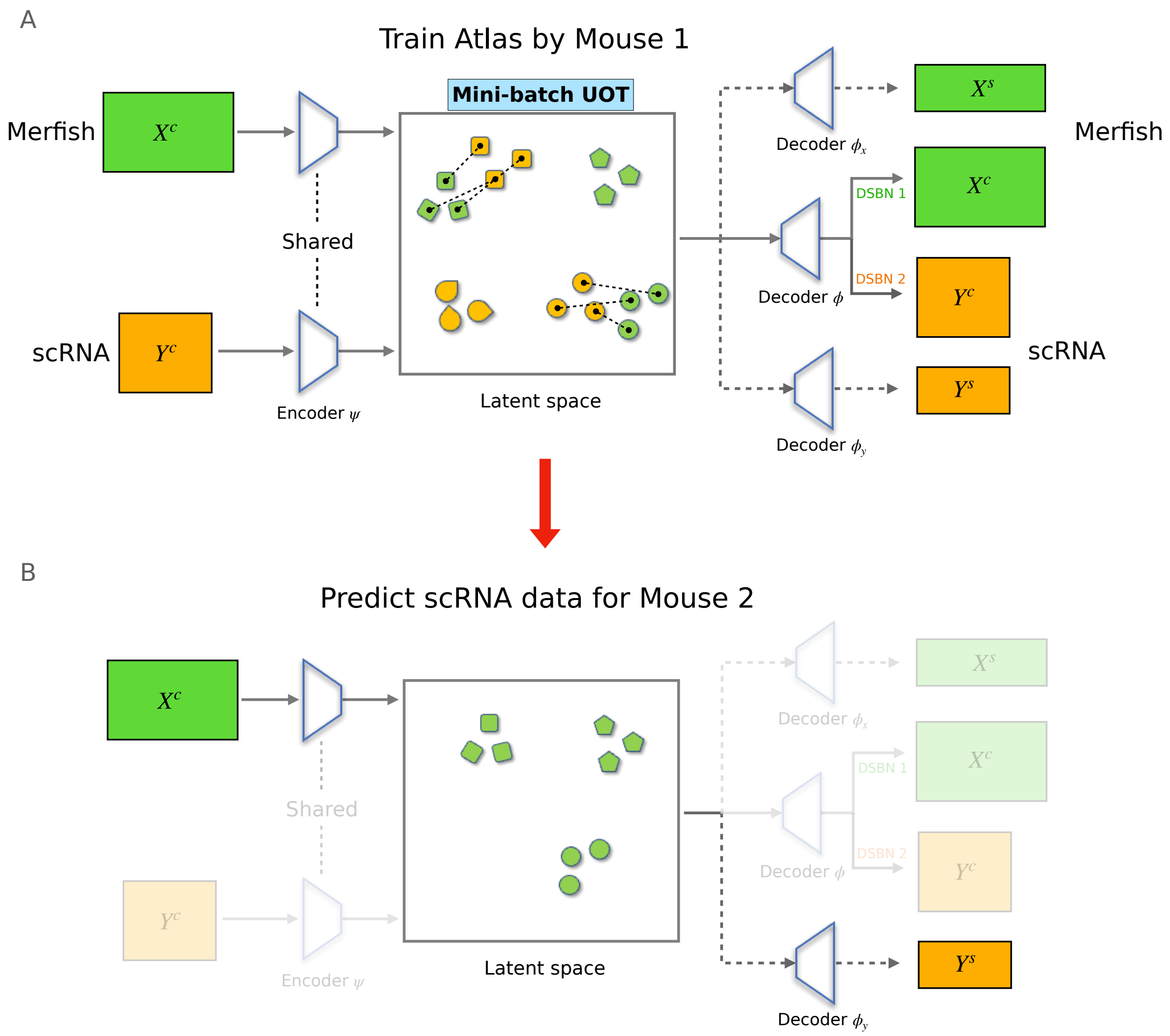


**Supplementary Figure 4. Online prediction. (A)** uniPort trained an atlas by scRNA and MERFISH data from mouse 1. **(B)** uniPort predicted scRNA through MERFISH from mouse 2.

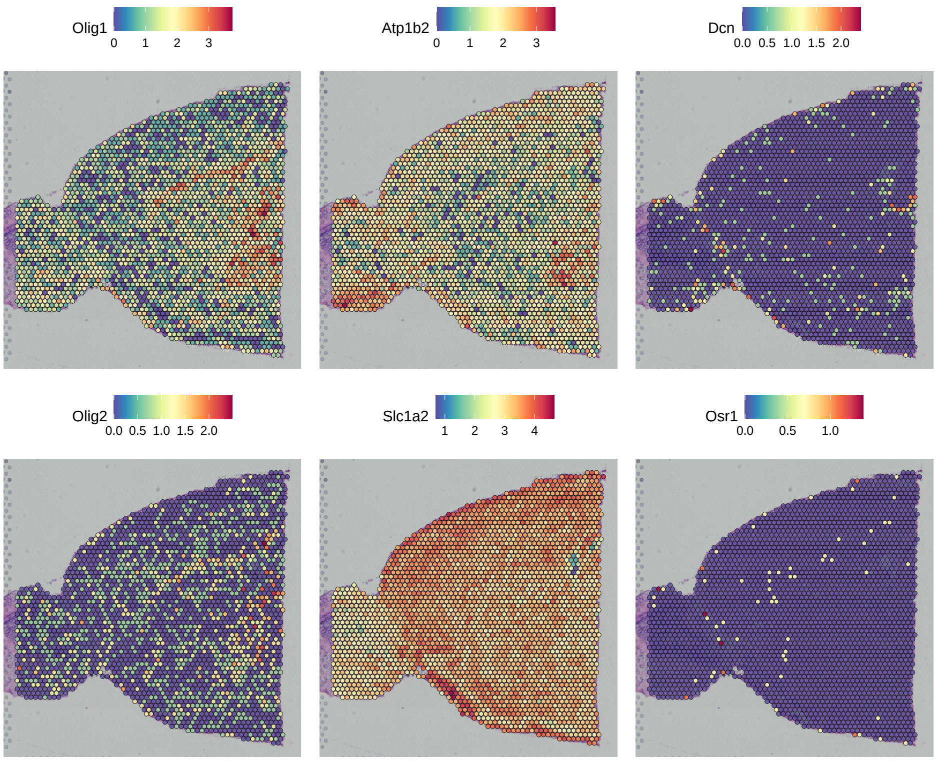


**Supplementary Figure 5. Marker genes in adult mouse brain ST data.** Left panel: Olig1 and Olig2 of cluster Oligo; Middle panel: Atp1b2 and Slc1a2 of cluster Astro; Right panel: Dcn and Osr1 of cluster VLMC.

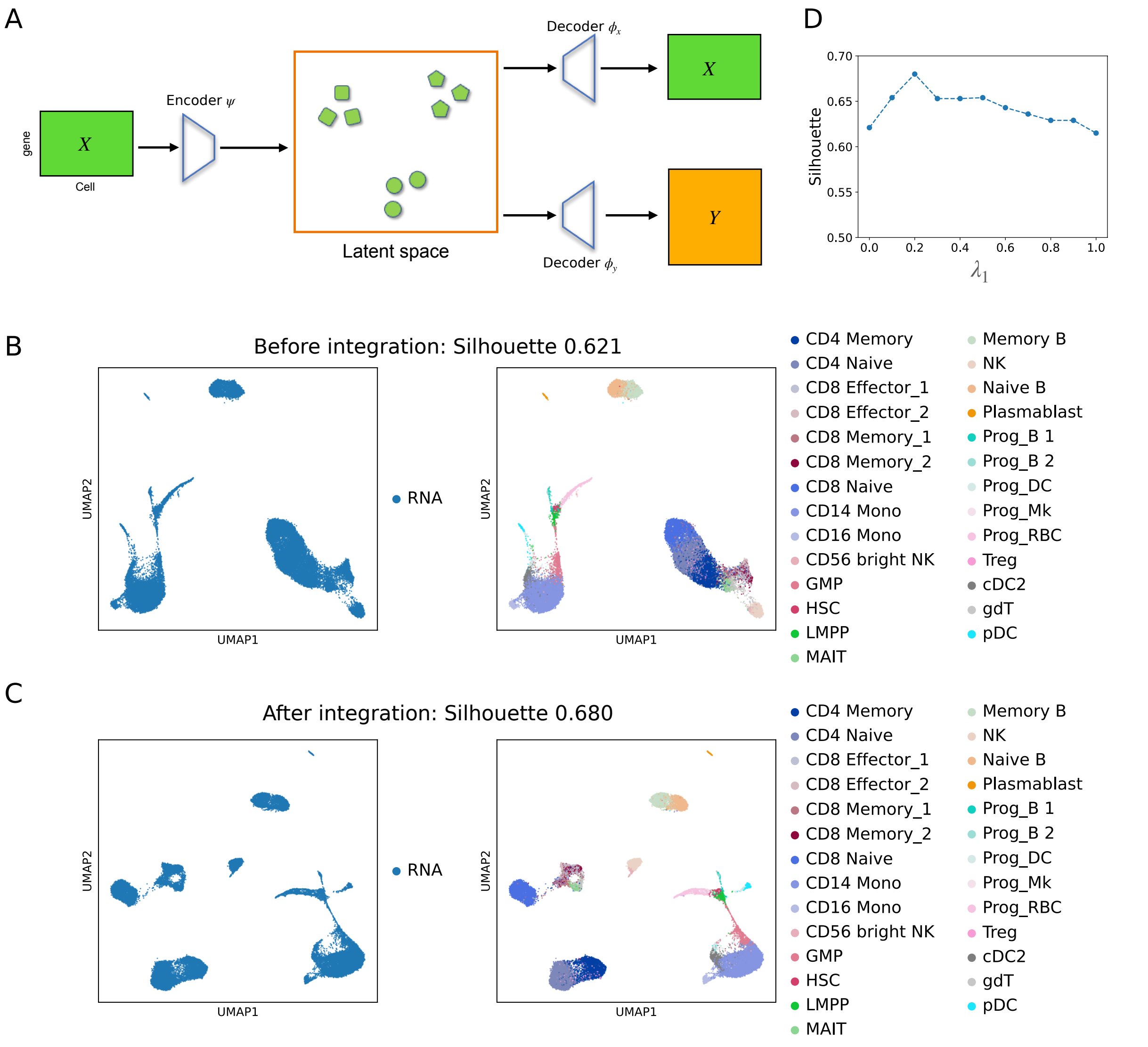


**Supplementary Figure 6. Vertical integration.** (**A**) uniPort_v takes one dataset as input and projects it into a low-dimensional latent space by a probabilistic decoder. Then uniPort_v reconstructs different datasets through its specific decoders from cell embeddings in the latent space. (**B**) UMAP visualization of scRNA data without integration of CITE data. (**C**) UMAP visualization of scRNA data with integration of CITE data. (**D**) The change of Silhouette score with different choices of $\lambda_1$ parameter.

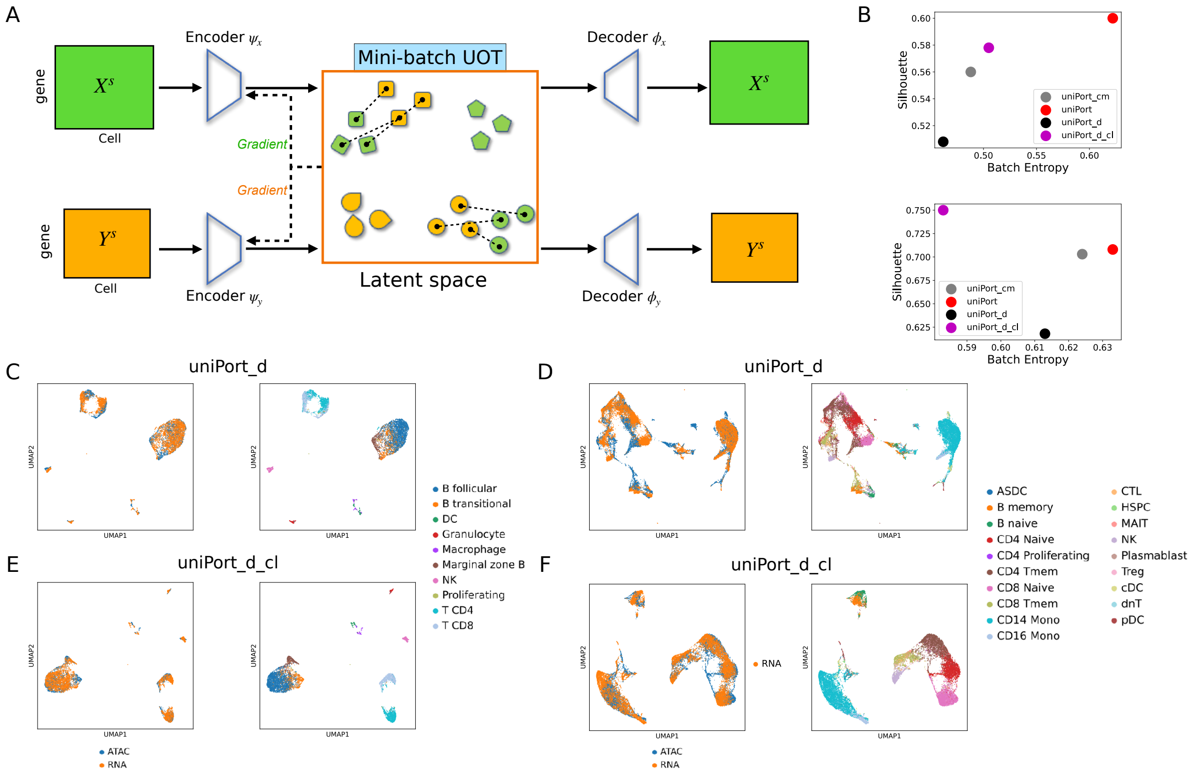


**Supplementary Figure 7. Diagonal integration. (A)** uniPort_d takes different datasets without aligned common genes as input, and projects them into a common cell-embedding space by different encoders. Then uniPort_d minimizes the Mini-batch UOT loss and reconstructs input through corresponding decoders. **(B)** Comparison of Batch Entropy and Silhouette scores of uniPort_d (uniPort without common genes), uniPort_cm (uniPort with only common genes), uniPort (uniPort with both common and specific genes) and uniPort_d_cl (uniPort_d with contrastive learning) in mouse spleen and PBMC data. **(C)** UMAP visualization of mouse spleen by uniPort_d without aligned common genes. **(D)** UMAP visualization of PBMC by uniPort_d without common genes. **(E)** UMAP visualization of mouse spleen by uniPort_d_cl with contrastive learning. **(F)** UMAP visualization of PBMC data by uniPort_d_cl with contrastive learning.

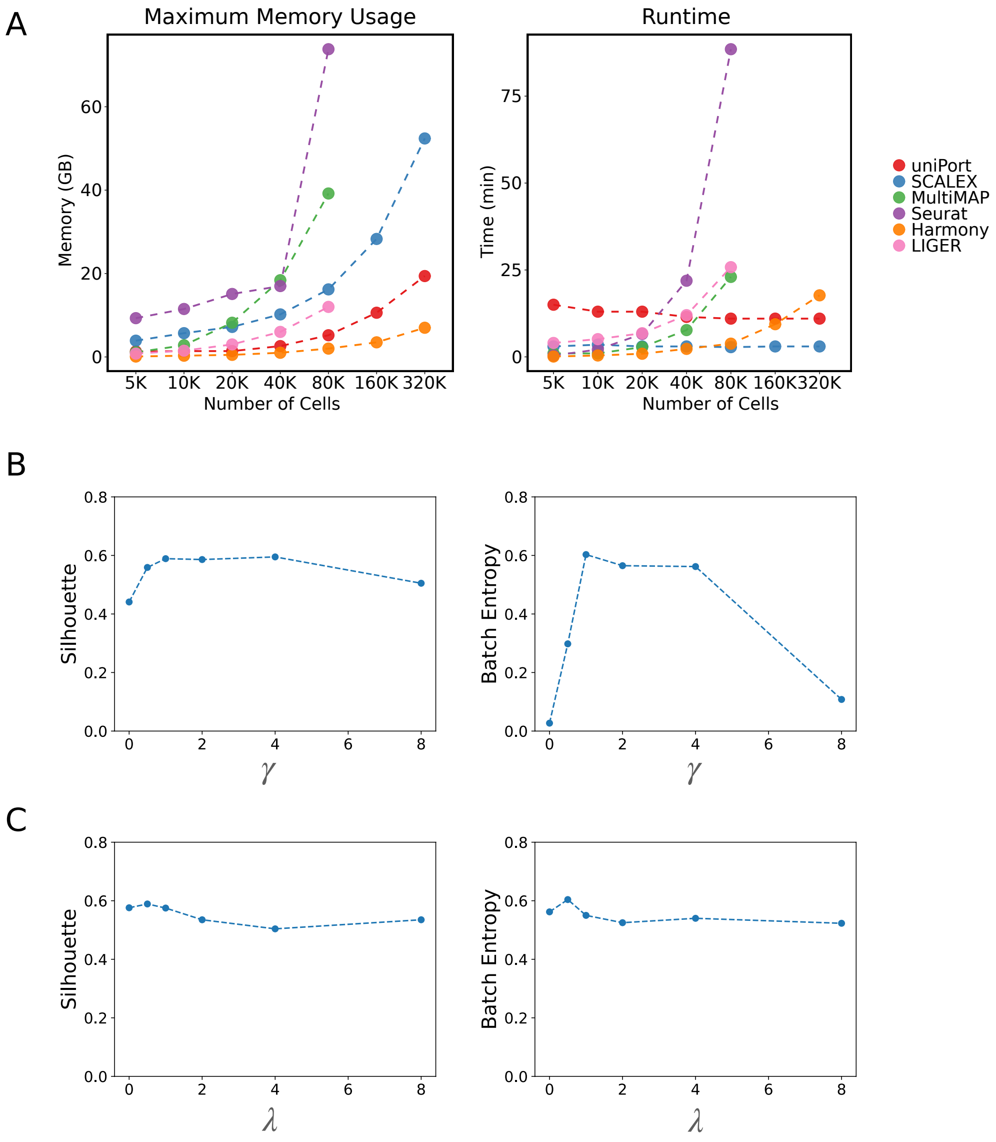


**Supplementary Figure 8. The computational cost and robustness of uniPort. (A)** Maximum memory usage and total runtime of different data sizes. **(B)** Changes of Silhouette and Batch Entropy scores with different choices of $\gamma$. **(C)** Changes of Silhouette and Batch Entropy scores with different choices of $\lambda$.

[2] Kai Cao, Yiguang Hong, and Lin Wan. Manifold alignment for heterogeneous single-cell multiomics data integration using Pamona. *Bioinformatics*, 08 2021.
